# Inhibition of Posterior Thalamic Nuclei Attenuates CGRP-induced Migraine-like Behavior in Mice

**DOI:** 10.1101/2025.03.03.641257

**Authors:** Agatha M. Greenway, Michael W. Huebner, Jayme S. Waite, Harold C. Flinn, Brandon J. Rea, Toby C. Buxton, Thomas L. Duong, Mengya Wang, Hailey L. Uemura, Nicholas O. Dorricott, Joseph O. Tutt, Kai Wang, Andrew F. Russo, Levi P. Sowers

**Author notes:** **Corresponding author:** L.P. Sowers. Department of Neurology, University of Utah, Salt Lake City, Utah, USA.

## Abstract

**Objective:** To determine whether induction of migraine-like symptoms in mice by calcitonin gene-related peptide (CGRP) requires activation of the posterior thalamic nuclei (PoT) in the brain.

**Background:** Previous research found that both optical activation of the PoT and injection of CGRP into the PoT are sufficient to induce light aversive behavior in mice. The PoT is well known as a sensory integration center of light and pain signals in the brain. However, whether this region is required for touch hypersensitivity and light aversion following peripheral administration of CGRP was not known.

**Methods:** The PoT was injected in two strains of mice, inbred C57BL/6J and outbred CD-1, with viral vectors expressing inhibitory chemogenetic Designer Receptors Exclusively Activated by Designer Drugs (DREADDs). The inhibitory DREADDs were activated by systemic intraperitoneal (ip) injection of two designer drugs, clozapine N-oxide (CNO) and DREADD agonist compound 21 (C21). We used ip injection of CGRP to induce migraine-like phenotypes and tested whether we could rescue these phenotypes by bilateral chemogenetic inhibition of the PoT. The light/dark assay was used to measure light aversive behavior (a surrogate for photophobia) and the plantar von Frey assay to measure hindpaw touch sensitivity (a surrogate for extracephalic allodynia).

**Results:** We successfully induced light aversive and hindpaw touch hypersensitivity phenotypes in mice using ip injections of CGRP. Activation of the inhibitory DREADDs in the PoT using ip CNO (5 mg/kg) was sufficient to partially rescue the touch hypersensitivity phenotype, but with off target effects in the control mice. Lowering the CNO dose to 1 mg/kg alleviated off target effects but was insufficient to rescue the touch hypersensitivity phenotype. On the other hand, C21 (1 mg/kg) fully rescued the touch hypersensitivity phenotype without any off target effects. Treatment with C21 also partially rescued the light aversion phenotype. These results were consistent across both C57BL/6J and CD-1 mouse strains.

**Conclusion:** Inhibition of the PoT fully rescues CGRP-induced touch hypersensitivity and partially rescues light aversion in mice, indicating that the PoT is necessary for touch hypersensitivity and partially necessary for light aversive behaviors. These data suggest the PoT is part of a central network that receives peripheral CGRP-induced signals and thus could be harnessed for future targeted therapeutics for migraine.

**Plain language summary:** The posterior thalamus is a central brain region that contributes to migraine pathophysiology when stimulated. In this study, we asked if inhibition of this brain region could alleviate migraine-like phenotypes in mice. We found that inhibition of the posterior thalamus fully rescues touch hypersensitivity and partially rescues light aversive behavior, suggesting that the posterior thalamus is necessary for migraine pathophysiology and could offer a potential therapeutic target for migraine.

## Introduction

Migraine is a complex, multifaceted neurological disorder, ranked globally as one of the leading causes of disability^1^. Migraine affects 15% of the US population, with a preponderance in females compared to males (approximately 2:1)^2^. In addition to head pain, migraine patients suffer from debilitating sensory abnormalities, including cutaneous allodynia (hypersensitivity to touch) and photophobia (hypersensitivity to light). Around 60% of people with migraine experience cutaneous allodynia^3^, with cephalic allodynia as the most common form. Approximately 25-50% of people with migraine experience extracephalic cutaneous allodynia^4–7^. Photophobia is reported by around 90% of migraine patients and is the most bothersome symptom after head pain^8, 9^.

Calcitonin gene-related peptide (CGRP) is a key neuropeptide in migraine pathogenesis and peripheral injection of CGRP can induce migraine-like phenotypes in both clinical and pre-clinical models^10–13^. However, it is not known how peripheral CGRP-induced migraine affects the central nervous system (CNS) and which brain regions are implicated in CGRP-induced migraine-like phenotypes. Here we use intraperitoneal (ip) CGRP injection as a translational model for acute migraine^14–16^. Over the past two decades, advances in migraine therapeutics have centered on the development of eight U.S. Food and Drug Administration-approved CGRP antagonist drugs, which are effective in about 50% of patients^17–22^.

Despite the success of CGRP-based drugs, there is a paucity of knowledge surrounding the central brain regions and mechanisms responsible for migraine pathophysiology. Specifically, how brain regions in the CNS contribute to peripherally administered migraine-inducing compounds like CGRP is an open question. Moreover, CGRP antagonist drugs are thought to mostly act in the periphery^21, 23–25^, so what role do central brain regions play in migraine phenotypes? Previous human imaging studies have defined brain regions contributing to migraine, including the thalamus^26–28^. Patients responsive to treatment with the CGRP monoclonal antibody erenumab demonstrate alterations in resting state functional connectivity, suggesting peripheral CGRP activity affects central connectivity^29^.

One key brain region implicated in migraine is the pulvinar region of the thalamus, which includes the posterior thalamic nuclei (PoT). The PoT is a collection of thalamic nuclei that are responsible for integrating and reciprocally relaying neural impulses between multiple sensory inputs and regions of the cerebral cortex^30–32^. Importantly, clinical and preclinical studies identified the PoT as a crucial region in migraine pathophysiology and this region has been linked to somatosensory pain signals^33–38^. The PoT receives direct inputs from the spinal trigeminal nucleus (SpV) via the trigeminovascular pathway^35, 39^. Dura-sensitive neurons in laminae I and V of the SpV that project to the PoT receive information from sensory neurons in the trigeminal ganglion, which innervate the head^35, 39, 40^. Exacerbation of headache by light is modulated at the thalamic level by non-image forming retinal ganglion cells converging on dura-sensitive neurons in several posterior thalamic nuclei^35^. These thalamic neurons then project to a variety of pain and light processing cortical regions, including the primary somatosensory cortex and primary and secondary visual cortices^35^. The combination of sensory inputs into the PoT could be responsible for both cutaneous allodynia and photophobia in migraine. Supporting this idea, we previously demonstrated that both direct injection of CGRP into the PoT and optical stimulation of the PoT were sufficient to induce migraine-like phenotypes in mice^37^. Together these data suggest the PoT is an important sensory integration center in migraine.

To determine if involvement of the PoT is necessary for migraine-like phenotypes, we employed a chemogenetic strategy using inhibitory designer receptors exclusively activated by designer drugs (DREADDs). Specifically, we used the modified form of the human M4 muscarinic receptor (hM4Di) which is activated by ip injection of either clozapine N-oxide (CNO) or DREADD agonist compound 21 (C21). We induced migraine-like states in mice using ip CGRP injection and investigated whether these states are rescued with DREADD inhibition of the bilateral PoT. We used light aversive behavior as a surrogate measure of photophobia and hindpaw touch hypersensitivity as a surrogate of extracephalic cutaneous allodynia. Unfortunately, cephalic allodynia could not be measured in both strains of mice since the periorbital region was hypersensitive following the surgical procedure. We have previously reported an issue with reproducibly habituating mice to von Frey filaments for measuring periorbital sensitivity^41^. We found that DREADD inhibition of the PoT fully rescues hindpaw touch hypersensitivity and partially rescues light aversion in mice. These data suggest the PoT is an important integration site for peripheral CGRP signals and is necessary for some migraine-like sensory phenotypes.

## Materials and methods

### Animals

A total of 105 wild type C57BL/6J mice (https://www.jax.org/strain/000664) at 10-13 weeks of age underwent surgery (51 females, 54 males), with female and male average starting body weights of 20 g and 26 g, respectively. A total of 30 wild type CD-1 mice (https://www.criver.com/products-services/find-model/cd-1r-igs-mouse?region=3611) at 12 weeks of age underwent surgery (19 females, 11 males), with female and male average starting body weights of 34 g and 43 g, respectively. We observed no behavior differences between sexes and therefore the data were analyzed together. All animals were housed in a temperature controlled-vivarium in groups of 2-5 per cage on a 12-hour light cycle with access to water and food *ad libitum*. Animal procedures were approved by the Iowa City Veterans Affairs (VA) and University of Iowa Institutional Animal Care and Use Committees (IACUC) and followed the ARRIVE (Animal Research: Reporting of *In Vivo* Experiments) guidelines and performed in accordance with the National Institutes of Health standards.

### Stereotactic injection of adeno-associated viruses (AAV) into the PoT

Stereotaxic surgery for bilateral virus injections into the PoT was performed under isoflurane anesthesia (induction 5%, maintenance 1.5-2%). All standard operating procedures defined by the VA and University of Iowa IACUC for rodent surgery were followed in preparation of the mouse. hM4Di was expressed via bilateral injection into the PoT using an adeno-associated virus 5 (AAV5) containing a fusion gene of hM4Di and an mCherry reporter, with expression driven by calmodulin kinase IIa (CaMKIIa) promoter (AAV5-CaMKIIa-hM4Di-mCherry), or the control vector (AAV5-CaMKIIa-mCherry), both obtained from Addgene. Coordinates for the PoT injections from bregma were: anterior/posterior (AP), −0.18 cm posterior; medial/lateral (ML), +/−0.11 cm lateral; and dorsal/ventral (DV), −0.32 cm from pial surface. The injection rate was 200 nL/minute for 1 minute (Micro4, Model UMP3 pump), using a 10 µL gas tight Hamilton syringe (Hamilton Company) and a 30-gauge needle. The needle was left in place for an additional 5 minutes post injection to reduce fluid tracking up the needle. Incisions were closed with coated VICRYL 7-0 suture (Ethicon, Inc). Analgesia (ip meloxicam, 0.2 mg/kg) was administered at the conclusion of injections and for 2 consecutive days post surgery.

### Activation of hM4Di designer receptor

We tested two designer drugs to activate hM4Di, CNO and C21 (HelloBio). The ligands were mixed in house with 1X phosphate buffered saline (PBS; Hyclone^TM^). CNO was tested at 2 different doses, 5 mg/kg and 1 mg/kg, and C21 was tested at 1 mg/kg^42, 43^.

### Plantar von Frey assay

The von Frey assay was used to test touch hypersensitivity in the plantar region of the right hindpaw as previously described^15^. The mice were habituated to the behavior room for 1 hour prior to testing. Data are shown as 50% thresholds (g) which were transformed into logarithmic format to allow for linear analysis and graphing of figures^44^.

### Light/dark assay

The twelve individual testing chambers consist of a plexiglass open field divided equally into two zones using a dark insert, as previously described^45, 46^. The light/dark and motility data were collected using Activity Monitor version 7.06 (Med Associate Inc), and all mice were tested in the chamber with LED lighting at 2.7 × 10^4^ lux^45^. Data were collected for a total of 30 minutes and analyzed in sequential 5 minute intervals. Time spent in the light zone per 5 minute interval was averaged to determine average time spent in light.

### Histology

To determine the presence and spread of viral injection, following conclusion of behavioral testing, mice were deeply anesthetized with ketamine/xylazine (87.5 mg/kg/12.5 mg/kg, ip), perfused transcardially using 1X PBS and 4% paraformaldehyde, cryoprotected using a sucrose gradient, and frozen in tissue freezing medium at -80 °C until use, following standard procedures^37, 47^. The brains were cut into 30-40 μm coronal slices and mounted onto Superfrost Plus slides (Fisher Scientific) using VECTASHIELD HardSet antifade mounting medium with DAPI (Vector Laboratories). Images were acquired using a light microscope (Olympus, VX61VS) equipped with an Hamamatsu C13440 camera with tetramethylrhodamine isothiocyanate (TRITC) filter and 20X magnification and processed using the INFINITY ANALYZE software (Lumenera Corporation).

### Experimental design

After surgeries, mice were given 3 weeks for sufficient recovery and viral expression prior to behavioral testing. Mice were baselined in the behavior assay 2 days prior to testing for acclimation (data not shown). On the day of testing, mice were allowed to acclimate to the behavior room for at least 1 hour prior to commencement of behavioral assays. On test day 1, the mice were divided evenly and randomized by sex and viral injection and given a 10 µl/g ip injection of either rat ⍺-CGRP (CGRP; Sigma-Aldrich) at 0.1 mg/kg (diluted in 1X PBS) or an equivalent volume of PBS. The mice were returned to their home cage for 30 minutes post ip injection prior to testing in the behavior assay. Two days later, the mice were tested again (test day 2), with the same treatment groups but receiving two ip injections^14, 48^. The first injection was the designer drug at 5 µl/g. 10 minutes after the designer drug injection^42, 49^, a second injection was administered consisting of PBS or CGRP at 5 µl/g. To maintain the same total injection volume per testing day (10 µl/g), the concentration of CGRP injection was doubled on test day 2 (0.2 mg/kg) and the volume of CGRP (and equivalent PBS) was halved (5 µl/g). This ensured total injection volume remained in accordance with standard procedures set by University of Iowa IACUC. 30 minutes after the second injection, mice were retested in the behavior assay. Mice were returned to their home cages between and following injections. Testing paradigms are shown for the plantar von Frey assay and the light/dark assay in respective figures. Mice were tested in the light/dark assay first and von Frey assay second, with at least 1 week between assays. To increase sample size whilst minimizing mouse numbers, we implemented a crossover experimental design. Following completion of the first assay, where the mice were either treated with CGRP or PBS, the respective treatment groups were switched and the mice were retested in the same assay at least 2 days later, following the same paradigm shown in the figure schematics below. Mice that were first tested with CGRP treatment, were retested in the same assay with PBS, and vice versa. During each experiment, the experimenter was blinded to genotypes and treatments. All behavioral testing was conducted between the hours of 8:00 am and 5:30 pm.

### Statistical analysis

A power analysis was performed prior to the study for sample size estimation based on previous studies from the laboratory using ClinCalc.com. An alpha of 0.05 and a power of 0.80 was used which determined that 10 mice in each group were needed per behavior assay. Data were graphed using GraphPad Prism 10 and analyzed in R, with statistical outputs reported in Table 1. Significance was set at *p* ≤ 0.05. To account for repeated measurements in the cross-over design due to paired groups, the random effect model was used. This model uses all data points including those data points that have no paired data points. When there are no repeated measures (no crossover; unpaired groups only), a linear regression model was used. Error bars represent ± standard error of mean (SEM). We define a partial rescue as no significant difference between designer drug + CGRP versus CGRP, and designer drug + CGRP versus designer drug + PBS. A full rescue is defined as a significant difference between designer drug + CGRP versus CGRP, and no significant difference between designer drug + CGRP versus designer drug + PBS.

**Table 1:** Statistical Analysis.

| Figure # | Analysis type and sample size |
| --- | --- |
| 1B | PBS (n = 5) vs CNO + PBS (n = 10): random effect model, p = 0.1858042<br>CGRP (n = 5) vs CNO + CGRP (n = 11): random effect model, p = 0.4472274<br>CNO + PBS (n = 10) vs CNO + CGRP (n = 11): random effect model, p = 0.6658471<br>PBS (n = 5) vs CGRP (n = 5): linear regression model, p = 0.0227702 |
| 1C | PBS (n = 5) vs CNO + PBS (n = 15): random effect model, p = 0.2267849<br>CGRP (n = 10) vs CNO + CGRP (n = 15): random effect model, p = 0.0589026<br>CNO + PBS (n = 15) vs CNO + CGRP (n = 15): random effect model, p = 0.0968951<br>PBS (n = 5) vs CGRP (n = 10): linear regression model, p = 0.025252 |
| 1D | All analyses used the random effect model<br>PBS (n = 7) vs CNO + PBS (n = 7): p = 0.6195286<br>CGRP (n = 6) vs CNO + CGRP (n = 6): p = 0.3739151<br>CNO + PBS (n = 7) vs CNO + CGRP (n = 6): p = 0.0103153<br>PBS (n = 7) vs CGRP (n = 6): p = 0.0260111 |
| 1E | All analyses used the random effect model<br>PBS (n = 8) vs CNO + PBS (n = 8): p = 0.9496605<br>CGRP (n = 8) vs CNO + CGRP (n = 8): p = 0.8812002<br>CNO + PBS (n = 8) vs CNO + CGRP (n = 8): p = 0.008655<br>PBS (n = 8) vs CGRP (n = 8): p = 0.0051219 |
| 2B | All analyses used the random effect model<br>PBS (n = 20) vs C21 + PBS (n = 20): p = 0.2459639<br>CGRP (n = 21) vs C21 + CGRP (n = 21): p = 0.6111491<br>C21 + PBS (n = 20) vs C21 + CGRP (n = 21): p = $8.9377243 \times 10^{-4}$<br>PBS (n = 20) vs CGRP (n = 21): p = $2.4267654 \times 10^{-4}$ |
| 2C | PBS (n = 22) vs C21 + PBS (n = 22): random effect model, p = 0.3316084<br>CGRP (n = 21) vs C21 + CGRP (n = 21): random effect model, p = 0.0036087<br>C21 + PBS (n = 22) vs C21 + CGRP (n = 21): random effect model, p = 0.0545357<br>PBS (n = 22) vs CGRP (n = 21): linear regression model, p = 0.0052022 |
| 2D | All analyses used the random effect model<br>PBS (n = 11) vs C21 + PBS (n = 11): p = 0.068424<br>CGRP (n = 11) vs C21 + CGRP (n = 11): p = 0.0480154<br>C21 + PBS (n = 11) vs C21 + CGRP (n = 11): p = 0.1360316<br>PBS (n = 11) vs CGRP (n = 11): p = 0.0227095 |
| 3B | All analyses used the random effect model<br>PBS (n = 13) vs C21 + PBS (n = 13): p = 0.0582875<br>CGRP (n = 15) vs C21 + CGRP (n = 15): p = 0.8552742<br>C21 + PBS (n = 13) vs C21 + CGRP (n = 15): p = 0.0958865<br>PBS (n = 13) vs CGRP (n = 15): p = 0.0476101 |
| 3C | All analyses used the random effect model<br>PBS (n = 12) vs C21 + PBS (n = 12): p = 0.7634183<br>CGRP (n = 14) vs C21 + CGRP (n = 14): p = 0.1140345<br>C21 + PBS (n = 12) vs C21 + CGRP (n = 14): p = 0.542061<br>PBS (n = 12) vs CGRP (n = 14): p = 0.0042095 |
| 3D | All analyses used the random effect model<br>PBS (n = 11) vs C21 + PBS (n = 11): p = 0.7756044<br>CGRP (n = 11) vs C21 + CGRP (n = 11): p = 0.1159398<br>C21 + PBS (n = 11) vs C21 + CGRP (n = 11): p = 0.1521276<br>PBS (n = 11) vs CGRP (n = 11): p = 0.039174 |
| Supp. 1A | All analyses used the random effect model<br>PBS (n = 7) vs C21 + PBS (n = 7): p = 0.1313264<br>CGRP (n = 6) vs C21 + CGRP (n = 6): p = 0.4721717<br>C21 + PBS (n = 7) vs C21 + CGRP (n = 6): p = 0.0505061<br>PBS (n = 7) vs CGRP (n = 6): p = 0.3151376 |
| Supp. 1B | All analyses used the random effect model<br>PBS (n = 5) vs C21 + PBS (n = 5): p = 0.1256103<br>CGRP (n = 7) vs C21 + CGRP (n = 7): p = 0.3997665<br>C21 + PBS (n = 5) vs C21 + CGRP (n = 7): p = 0.1002211<br>PBS (n = 5) vs CGRP (n = 7): p = 0.0015169 |
| Supp. 1C | All analyses used the random effect model<br>PBS (n = 4) vs C21 + PBS (n = 4): p = 0.1178043<br>CGRP (n = 4) vs C21 + CGRP (n = 4): p = 0.2867029<br>C21 + PBS (n = 4) vs C21 + CGRP (n = 4): p = 0.5461774<br>PBS (n = 4) vs CGRP (n = 4): p = 0.1526172 |

A total of 19 C57BL/6J mice and 5 CD-1 mice were excluded during testing due to either injection complications, recording issues or cage flooding. Due to viral injection mistargeting, an additional 19 C57BL/6J and 8 CD-1 mice were excluded; these are graphed separately in the supplemental figures.

## Results

### Inhibition of the PoT using CNO partially rescues CGRP-induced touch hypersensitivity in C57BL/6J hM4Di mice, but with off target effects

Approximately 60% of patients with migraine experience cutaneous allodynia during an attack ^3^. Cephalic allodynia is the more common form of cutaneous allodynia, but 25-50% of people with migraine report extracephalic allodynia^4–7^. Unfortunately, cephalic allodynia could not be measured in mice since the periorbital region was hypersensitive following the surgical procedures, as aforementioned. Thus, we focused on extracephalic allodynia by measuring touch hypersensitivity in the plantar region of the hindpaw in mice. We previously demonstrated that ip CGRP injection induces hindpaw touch hypersensitivity and light aversion in mice^14, 15^. Moreover, CGRP injection into, and optical stimulation of, the PoT induced migraine-like behaviors^37^ suggesting the PoT is sufficient to induce migraine-like behaviors in mice. Therefore, we tested if the PoT was necessary for CGRP-induced hindpaw touch sensitivity using CNO-mediated DREADDs (5 mg/kg). We performed bilateral inhibitory DREADD viral injections into the PoT of mice and tested if PoT inhibition can rescue CGRP-induced touch hypersensitivity. On test day 1, the mice were injected with ip CGRP (or PBS control) to induce a migraine-like state^10, 11^ before von Frey testing (Figure 1A left panel). Two days after test day 1, the mice received two ip injections of CNO + CGRP (or CNO + PBS) before being tested again in the von Frey assay (Figure 1A right panel, test day 2). CNO was administered 10 minutes before CGRP to activate hM4Di^49^. Previous studies from our group showed that repeated injections of CGRP do not affect behavior in these assays when two days are given between injections^14, 48^, and repeated injections during these experiments did not change behavioral results (data not shown).

**Figure 1:**
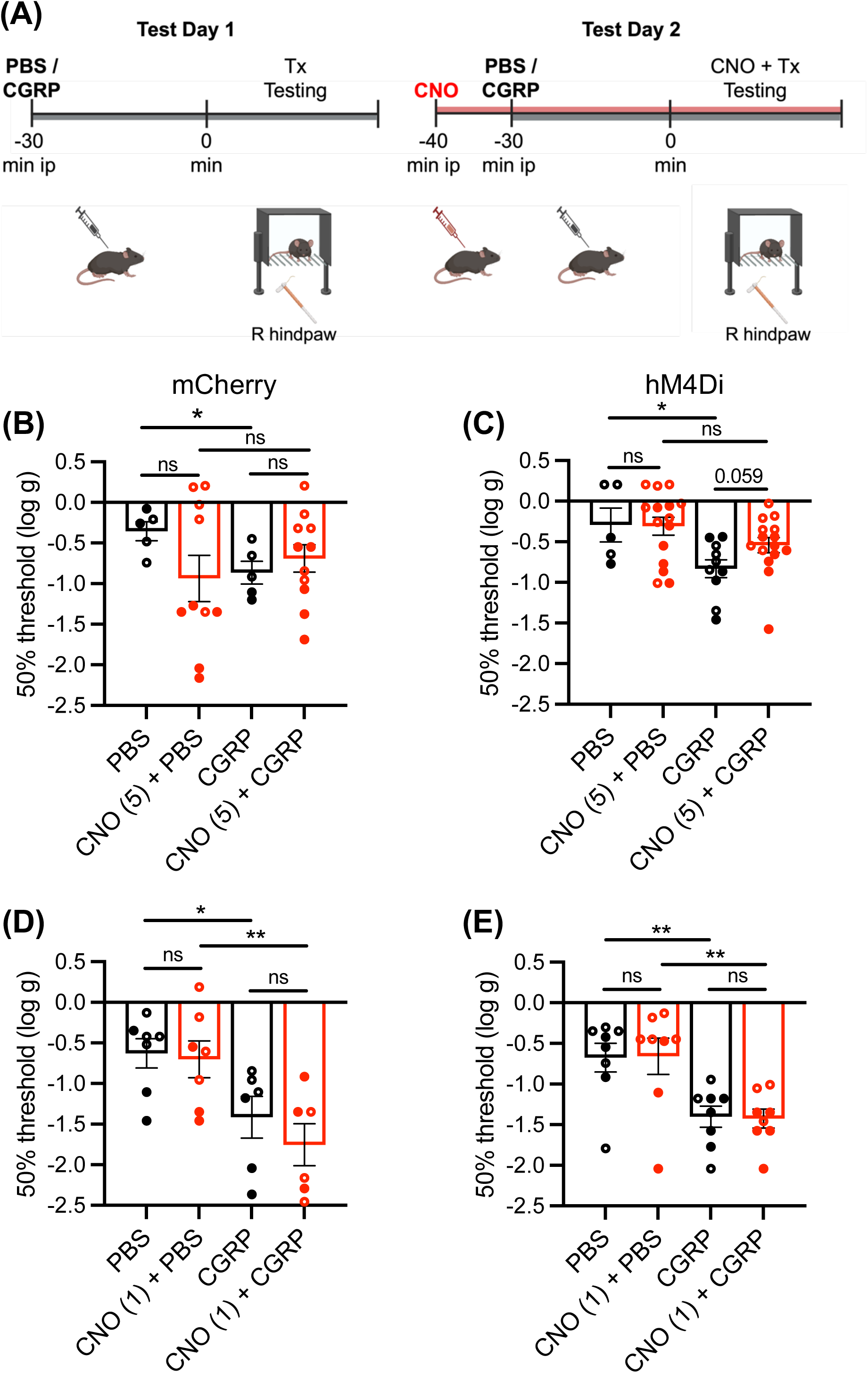
DREADD inhibition of the PoT with CNO partially rescues CGRP-induced touch hypersensitivity in C57BL/6J hM4Di mice with off target effects. (**A**) schematic demonstrating testing procedure for assessing plantar touch hypersensitivity in the right hindpaw. Tx = treatment (PBS/CGRP). (**B** and **C**) mCherry control mice (B) and hM4Di experimental (C) C57BL/6J mice receiving 1X PBS or CGRP (0.1 mg/kg) treatment, and CNO (5 mg/kg) + PBS or CNO + CGRP treatment. (**B**) CNO + PBS administration trends toward a decrease in withdrawal threshold compared to PBS treatment. CGRP treatment in mCherry mice induces a significant decrease in withdrawal threshold compared to PBS, which is not rescued by CNO administration. (**C**) PBS treatment in hM4Di mice and CNO + PBS treatment show no significant difference in withdrawal threshold. CGRP treatment induces a significant decrease in withdrawal threshold in hM4Di mice compared to PBS, and CNO + CGRP trends towards a rescue in withdrawal threshold. (**D** and **E**) mCherry control mice (D) and hM4Di experimental (E) C57BL/6J mice receiving 1X PBS or CGRP (0.1 mg/kg) treatment, and CNO (1 mg/kg) + PBS or CNO + CGRP treatment. (**D**) PBS treatment in mCherry mice and CNO + PBS treatment exhibit no significant difference in withdrawal threshold. CGRP treatment in mCherry mice induces a significant decrease in withdrawal threshold, which is not rescued by CNO administration. (**E**) PBS and CNO + PBS treatments in hM4Di mice show no significant difference in withdrawal threshold. CGRP treatment induces a significant decrease in withdrawal threshold for hM4Di mice compared to PBS treatment, and CNO + CGRP does not rescue this. CNO + CGRP treatment withdrawal threshold is significantly lower than CNO + PBS. Open and closed symbols denote males and females, respectively. Data were transformed to enable statistical comparisons to be made. Error bars: +/-standard error of the mean (SEM). \**p* ≤ 0.05. (**B - E**) 1 independent cohort of C57BL/6J mice with crossover design. For exact p-values and sample sizes, see Table 1.

C57BL/6J mice expressing the mCherry (control) viral vector that received an ip CGRP injection showed a significant decrease in the right hindpaw withdrawal threshold compared to mice receiving ip PBS (Figure 1B). Although there is no statistical difference between the PBS and CNO + PBS treatment conditions, there is a decreasing trend in withdrawal threshold for the CNO + PBS treatment compared to the PBS treatment condition, suggesting an off target effect of CNO. No significant difference was observed between the CGRP and CNO + CGRP treatment conditions.

C57BL/6J mice expressing the hM4Di (DREADDs) viral vector that received an ip CGRP injection showed a significant decrease in the right hindpaw withdrawal threshold compared to mice receiving ip PBS (Figure 1C). When comparing CGRP to CNO + CGRP treatment we see a trend towards significance (*p* = 0.059) and no significant difference between CNO + CGRP versus CNO + PBS treatment. These results indicate a partial rescue of CGRP-induced touch hypersensitivity upon CNO-mediated inhibition of the PoT (see statistical analysis section for definition of partial versus full rescue). It is important to note that according to our power analysis, we are underpowered for the mCherry and hM4Di PBS and CGRP treatments (< 10 mice per treatment). However, this experiment was exploratory to identify whether CNO is the most suitable designer drug for the plantar von Frey assay.

We tested a lower dose of CNO (1 mg/kg) in an attempt to alleviate the off target effects we observed in the mCherry mice with 5 mg/kg CNO (Figure 1D and 1E). Whilst we successfully alleviated the off target effect (Figure 1D), the partial rescue was lost in the hM4Di CNO + CGRP treated mice, demonstrated by a significant difference between CNO + CGRP and CNO + PBS (Figure 1E).

### Inhibition of the PoT using C21 fully rescues CGRP-induced touch hypersensitivity in C57BL/6J and CD-1 hM4Di mice

Due to the observed off target effects with the use of CNO, we used an alternative designer drug in the plantar von Frey assay, C21, which unlike CNO, does not retroconvert to biologically active compounds^42^. Testing followed the same paradigm as previously described, but with C21 (1 mg/kg) as the designer drug (Figure 2A). In the mCherry C57BL/6J mice, there is no significant difference between PBS and C21 + PBS treatments, suggesting no off target effect of C21 administration (Figure 2B). We observe a significant decrease in withdrawal threshold in the CGRP treated mice compared to PBS treatment. Compared to CGRP treatment, there is no significant difference in withdrawal threshold with C21 + CGRP treatment. We also observe a statistically significant decrease in withdrawal threshold for C21 + CGRP compared to C21 + PBS treatment, which is consistent with a CGRP-induced touch hypersensitivity phenotype with no rescue as these mice lack the hM4Di receptor.

**Figure 2:**
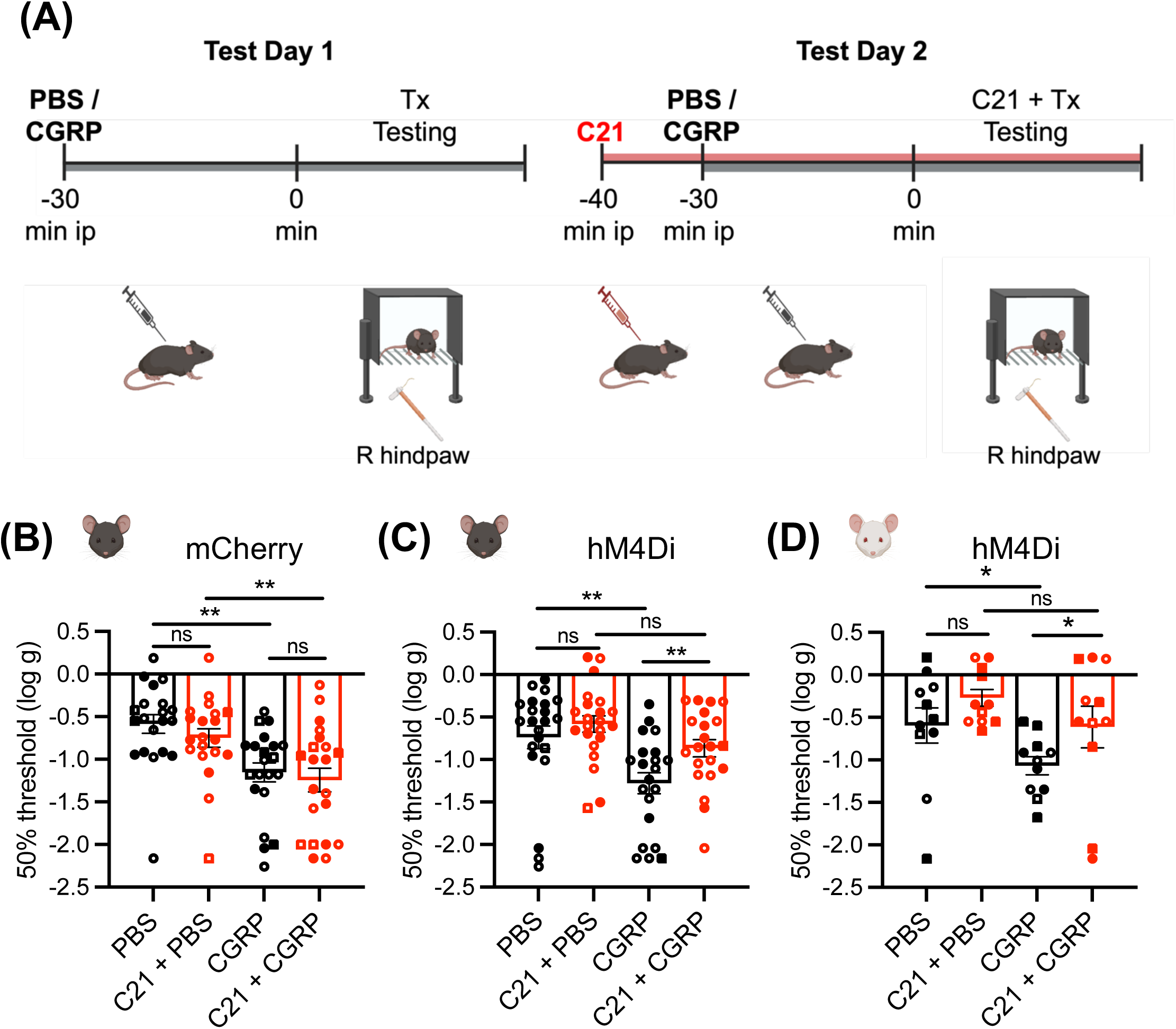
DREADD inhibition of the PoT using C21 fully rescues CGRP-induced touch hypersensitivity in both C57BL/6J and CD-1 hM4Di mice. (**A**) schematic demonstrating testing procedure for assessing plantar touch hypersensitivity in the right hindpaw. (**B** and **C**) mCherry control (B) and hM4Di experimental C57BL/6J mice (C), receiving 1X PBS or CGRP (0.1 mg/kg) treatment, and C21 (1 mg/kg) + PBS or C21 + CGRP treatment. (**B**) PBS treatment in mCherry mice and C21 + PBS treatment show no significant difference in withdrawal threshold. CGRP treatment in mCherry mice induces a significant decrease in withdrawal threshold compared to PBS, which is not rescued by C21 + CGRP administration. (**C**) PBS treatment in hM4Di mice and C21 + PBS treatment exhibit no significant difference in withdrawal threshold. CGRP treatment induces a significant decrease in hM4Di mice compared to PBS, and C21 + CGRP treatment significantly increases withdrawal threshold compared to CGRP treatment alone. (**D)** hM4Di experimental CD-1 mice receiving 1X PBS or CGRP (0.1 mg/kg) treatment, and C21 (1 mg/kg) + PBS or C21 + CGRP treatment. PBS treatment and C21 + PBS treatment in hM4Di mice show no significant difference in withdrawal threshold. CGRP treatment induces a significant decrease in withdrawal threshold hM4Di mice compared to PBS treatment. C21 + CGRP significantly increases withdrawal threshold compared to CGRP alone. Open and closed symbols denote males and females, respectively. Square symbols denote mice that did not undergo histology due to technical difficulties. Data were transformed to enable statistical comparisons to be made. Error bars are mean +/- SEM. \**p* ≤ 0.05, \*\**p* ≤ 0.01. (**B** and **C**) 2 independent cohorts of C57BL/6J mice. (**D**) 1 independent cohort of CD-1 mice. For exact p-values and sample sizes, see Table 1.

In the C57BL/6J mice that express the hM4Di receptor, CGRP treatment induces a significant decrease in withdrawal threshold compared to PBS, indicating a CGRP-induced touch hypersensitivity phenotype (Figure 2C). C21 + PBS treatment induces no significant difference in withdrawal threshold compared to PBS alone. C21 + CGRP treatment significantly increases withdrawal threshold compared to CGRP alone, indicating a full rescue of touch hypersensitivity upon inhibition of the PoT. Consistent with this finding, we see no statistical difference between C21 + CGRP treatment and C21 + PBS treatment.

We explored whether these results are consistent across different mouse strains and repeated the experiment with a CD-1 outbred strain of mice injected with experimental DREADDs virus (Figure 2D). In the hM4Di mice, we observed a significant CGRP touch hypersensitivity response compared to PBS, which was fully rescued by C21 + CGRP treatment and is consistent with the observed full rescue in C57BL/6J mice. For unknown reasons, the mCherry control CD-1 mice did not show a CGRP-induced decrease in withdrawal threshold compared to PBS (data not shown). Therefore, since the positive control was unsuccessful with this cohort, no conclusions can be drawn from the mCherry mice. Nonetheless, taken together, these data demonstrate that the PoT is necessary (full rescue) for the touch hypersensitivity phenotype in both C57BL/6J and CD-1 mice.

### Inhibition of the PoT using C21 partially rescues CGRP-induced light aversion in C57BL/6J and CD-1 hM4Di mice

Activation of the PoT via optical stimulation and CGRP injection demonstrated sufficiency to induce light aversive behavior in mice^37^. We wanted to determine whether the PoT is necessary for this phenotype, via DREADD-mediated inhibition. We decided to proceed with C21 (1 mg/kg) as the designer drug for this assay, as it showed no off target effects in the plantar von Frey assay. We induced a migraine-like state using ip 0.1 mg/kg CGRP (or PBS control) and exposed the mice to the light/dark assay for 30 minutes. Two days later, we reexposed the mice to the light/dark assay following a co-injection of C21 and CGRP or PBS control (Figure 3A).

**Figure 3:**
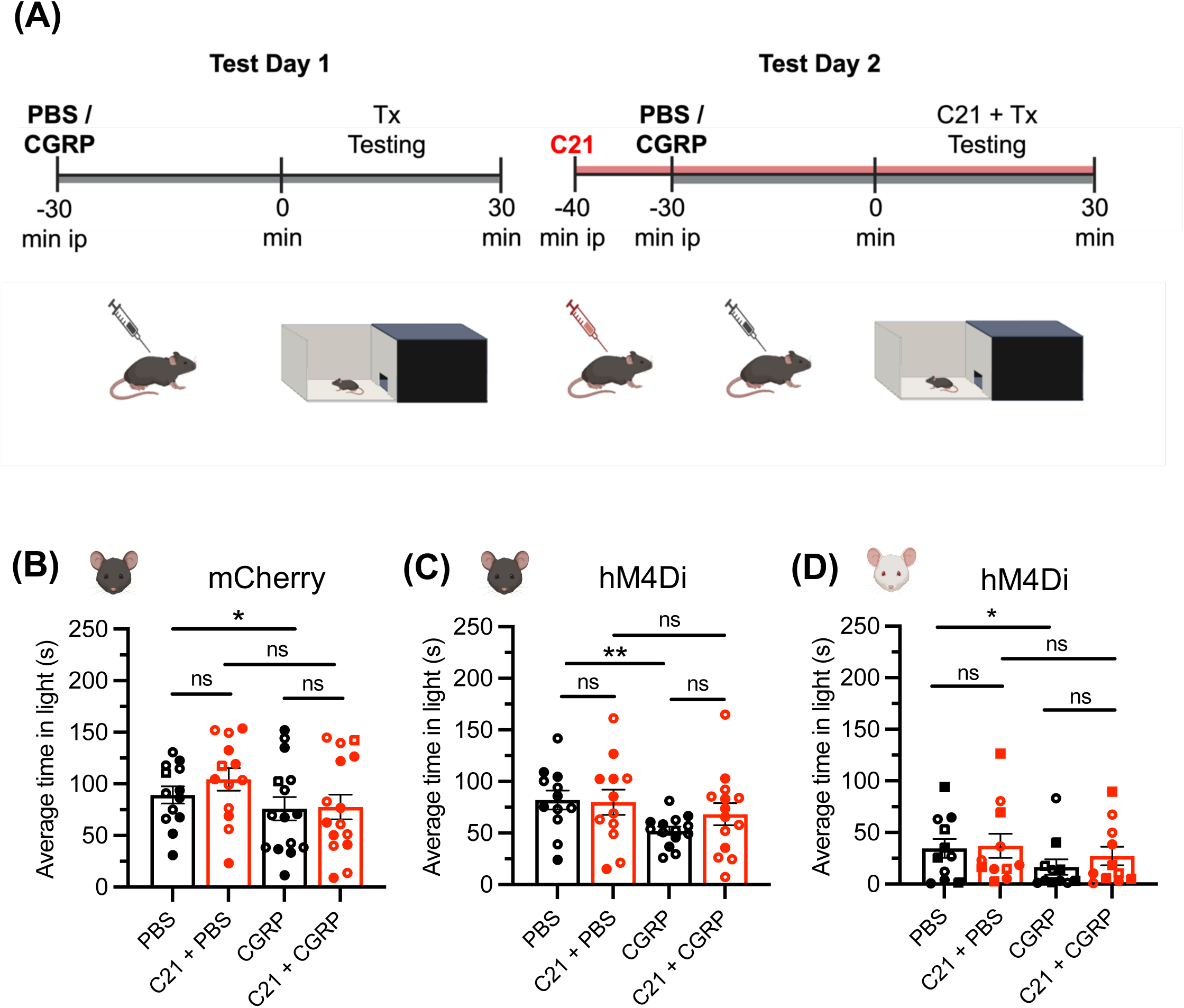
DREADD inhibition of the PoT partially rescues CGRP-induced light aversion in C57BL/6J and CD-1 hM4Di mice. (**A**) Schematic demonstrating testing strategy for light/dark assay. Data were collected in 5-minute intervals over the 30-minute testing period. (**B** and **C**) mCherry control C57BL/6J mice (B) and hM4Di experimental C57BL/6J mice (C) receiving 1X PBS or CGRP (0.1 mg/kg) treatment, and C21 (1 mg/kg) + PBS or C21 + CGRP treatment. (**B**) PBS treatment and C21 + PBS treatment in mCherry mice exhibit no significant difference in average time in light. CGRP treatment in mCherry mice induces a significant decrease in average time in light compared to PBS treatment. C21 + CGRP administration shows no significant decrease in time in light compared to CGRP treatment alone. (**C**) PBS treatment and C21 + PBS treatment exhibit no significant difference in average time in light in hM4Di mice. CGRP treatment in hM4Di mice induces a significant decrease in average time in light compared to PBS treatment. C21 + CGRP treatments induces a trend towards an increase in average time in light compared to CGRP treatment alone. (**D**) hM4Di experimental CD-1 mice receiving 1X PBS or CGRP (0.1 mg/kg) treatment and C21 (1 mg/kg) + PBS or C21 + CGRP treatment. PBS and C21 + PBS administration exhibit no significant difference in average time in light in hM4Di mice. CGRP administration causes a significant decrease in average time in light compared to PBS treatment and C21 + CGRP average time in light is not significantly different compared to CGRP alone or C21 + PBS. Open and closed symbols denote males and females, respectively. Square symbols denote mice that did not undergo histology due to technical difficulties. Average time in light +/- SEM shown. \**p* ≤ 0.05, \*\**p* ≤ 0.01. (**B** and **C**) 1 independent cohort of C57BL/6J mice. (**D**) 1 independent cohort of CD-1 mice. For exact p-values and sample sizes, see Table 1.

C57BL/6J mice expressing the mCherry virus exhibited a significant decrease in average time in light after ip CGRP administration compared to PBS (Figure 3B). This suggests a light aversive phenotype in the CGRP-induced migraine-like state. Upon C21 + PBS injection, there is no difference compared to PBS alone, suggesting no off target effect of C21 administration. There is also no difference between C21 + CGRP compared to CGRP alone. When comparing C21 + CGRP to C21 + PBS, we observe no significant difference.

C57BL/6J mice expressing the hM4Di receptor also exhibit a significant decrease in average time in light upon ip CGRP injection compared to PBS, indicating a light aversive phenotype (Figure 3C). There is no difference between C21 + PBS treatment and PBS alone. We also observe no statistical difference between CGRP and C21 + CGRP treatment, nor between C21 + CGRP treatment and C21 + PBS treatment, indicating a partial rescue of the light aversive phenotype.

CD-1 mice expressing the hM4Di receptor were then tested (Figure 3D). We observed a significant decrease in average time in light upon CGRP treatment compared to PBS. Treatment with C21 partially rescued the CGRP effect since was no significant difference between C21 + CGRP with either CGRP or PBS treated mice. As with the von Frey experiments, the same mCherry control CD-1 mice did not show a response to CGRP in the light aversion assay (data not shown). Therefore, since the positive control was unsuccessful with this cohort, no conclusions can be drawn from the mCherry mice. Despite this, these data demonstrate that the PoT is partially necessary (partial rescue) for the light aversive phenotype in both C57BL/6J and CD-1 mice.

### Identification of AAV expression in the bilateral PoT

This study involved a total of 4 independent cohorts of mice – 3 cohorts of C57BL/6J mice and 1 cohort of CD-1 mice. Histology was conducted on 2 of the C57BL/6J cohorts and the single CD-1 cohort. We were unable to conduct histological processing on the third C57BL/6J cohort due to complications in the post-testing pipeline. For the cohorts with histology conducted, the percentages of successful unilateral and bilateral hits in the PoT were determined and only mice with correctly targeted bilateral expression were included in the behavioral analyses. The 2 C57BL/6J cohorts combined had a bilateral (expression on both sides) PoT hit success rate of 70%, and a combined unilateral and bilateral injection hit success rate of 80%. Figure 4 demonstrates an example successful bilateral hit. For the CD-1 cohort, the bilateral PoT hit success rate was 68%, and the combined unilateral and bilateral injection hit success rate was 72%. The rostral border of detectable signal for fluorescent viral expression was -0.23 cm from bregma and the caudal most border was -3.27 cm from bregma. Given the high success rate for individual viral injections, the mice that did not have histology conducted were assumed to be bilateral hits in the PoT.

**Figure 4:**
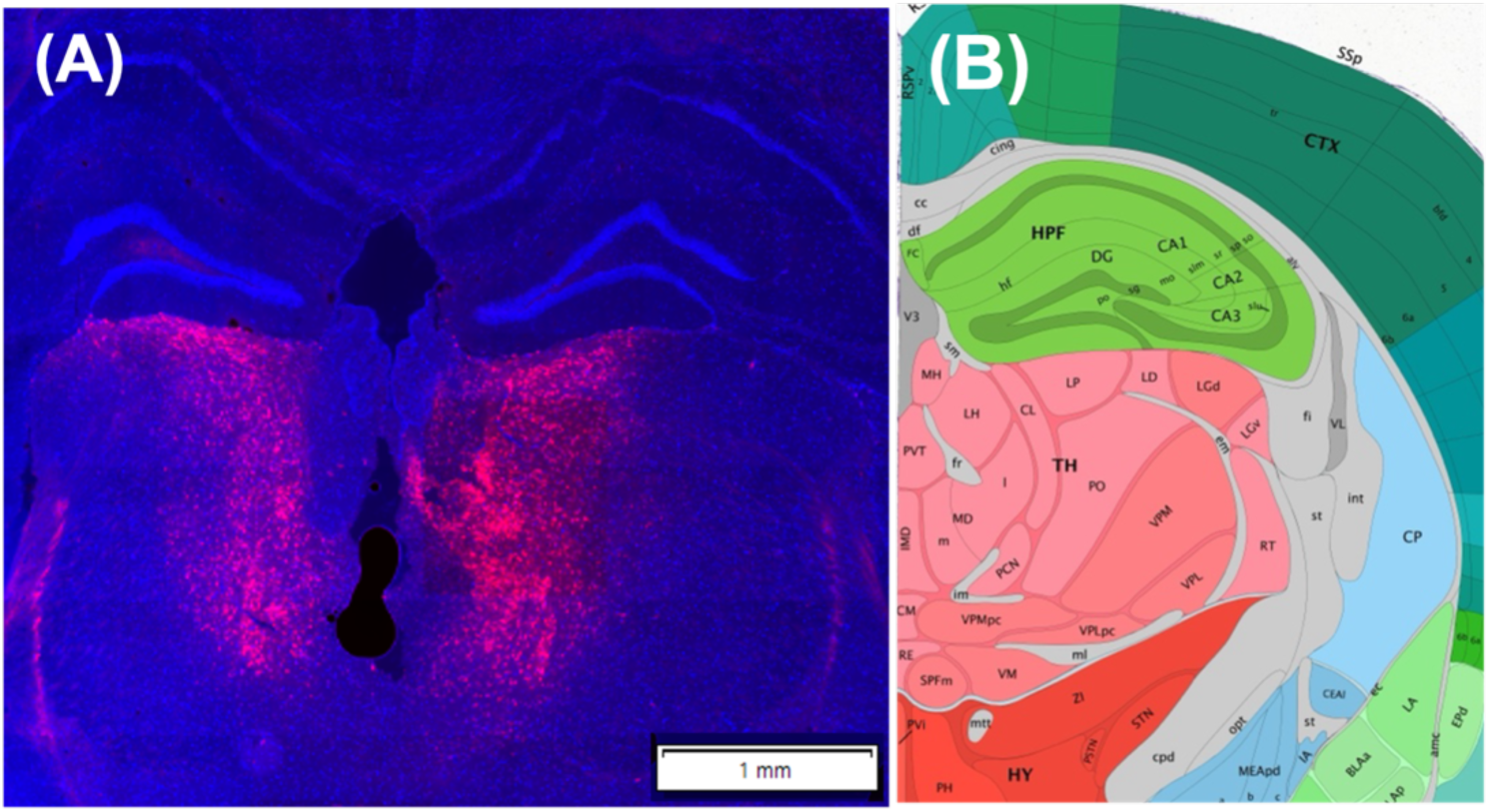
Validation of viral expression in the posterior thalamic nuclei. (**A**) Representative example of C57BL/6J mouse expression of AAV5-CaMKIIa-hM4Di-mCherry (red) in the posterior thalamic nuclei, counter stained with DAPI (blue). Scale bar 1mm shown. Magnification: 20x. (**B**) Anatomical annotations from the Allen Mouse Brain Atlas and Allen Reference Atlas, at the same slice position as A (image 74/132).

A mistargeted injection was defined as one that has no bilateral expression in the PoT, including unilateral expression, and the mistargeted von Frey data for both strains is graphed in Supplemental Figure 1. The light/dark assay data for the mistargeted mice are not shown due to the total sample size being very low (n ≤ 4). We acknowledge that we are underpowered in the number of mistargets to draw significant conclusions from these data. However, CGRP treatment induces a decrease in withdrawal threshold compared to PBS for hM4Di mice (Supplemental Figure 1A). Importantly, C21 activation of the mistargeted hM4Di receptor in the DREADDs mice fails to rescue the CGRP-induced touch hypersensitivity (Supplemental Figure 1B). These results are similar in the CD-1 mice (Supplemental Figure 1C). Taken together, these mistargeted data suggest the importance of bilateral inhibition of the PoT for effects on touch hypersensitivity.

## Discussion

Our previous study demonstrated that both optical activation and CGRP injection into the PoT is sufficient to induce light aversion in mice^37^. This suggested that the PoT may be a center for migraine-like hypersenstivity and that inhibition of this brain region may rescue migraine-like phenotypes. Since migraine can be initiated by peripheral administration of CGRP in humans^50^, we tested the role of the PoT following peripheral CGRP administration in mice. Here we show chemogenetic inhibition of the PoT rescued CGRP-induced touch hypersensitivity and partially rescued light aversion in two strains of mice.

Resting state human functional magnetic resonance imaging (fMRI) studies have highlighted that connectivity pathways between the PoT and cortical and subcortical regions involved in multi-dimensional pain processing are disrupted during migraine attacks^51^. Several studies have identified multiple neuronal pathways centered in the PoT that could mediate migraine-like allodynia and photophobia^32, 35, 39, 47, 52–54^. Light-sensitive PoT neurons project to a broad array of cortical sites, including multiple somatosensory regions, motor cortices, parietal association cortex, primary and secondary visual cortices, and insular cortices, with axonal projections spanning across cortical layers I-V^35, 52^. Dura and light sensitive projections from the PoT extend to the visual, auditory, retrosplenial agranular, motor and parietal association cortices^35^.

Extracranial allodynia has been linked to thalamic fMRI signals in response to mechanical and thermal stimulation^54^ and preclinical studies demonstrate that neurons in the posterior nucleus and ventral posteromedial nucleus of the thalamus are activated by mechanical and thermal stimulation in rats^39, 54^. These data highlight the PoT as a vital sensory integration center primed to regulate migraine hypersensitivity to light and touch, and our data support this.

The field of chemogenetics involving the use of DREADDs is rapidly advancing, with designer drug alternatives being developed with higher specificity, brain penetration and lower off target effects^55^. A necessary requirement of designer drug functionality is that it must be biologically inert and not possess pharmacological activity at non-DREADDs targets *in vivo*^43^. In the case of the first generation designer drug CNO, it reverse metabolizes back to its parent compound, clozapine, an antipsychotic that acts on a variety of endogenous targets to generate behavioral and physiological effects^43^. Indeed, it has been shown that clozapine, not CNO, crosses the blood brain barrier in mice to activate DREADDs^56^ and therefore suggests this conversion is necessary for DREADDs activation. Our results in the von Frey assay using CNO (5 mg/kg) appear to support this. However, this apparent off target effect is not present in the DREADD-expressing mice exposed to the same PBS + CNO treatment condition. It could be speculated that on target inhibition of the PoT was able to negate the off target effect of CNO in these mice. We lowered the concentration of CNO to see if the off target effects could be eliminated.

Unfortunately, although we eliminated the off target effects, we also no longer observed a rescue. C21 does not undergo back metabolism to clozapine^42, 57^, has high blood brain barrier penetrability and limited off target effects in the dose range of 0.4 -1 mg/kg^57^. We observed limited off target effects using C21 at 1 mg/kg, and recommend future behavior studies utilize C21, especially for use in von Frey experiments.

In the von Frey assay, we observe a CGRP-induced touch hypersensitivity phenotype. This is consistent with prior studies suggesting the proalgesic actions of peripheral CGRP on cutaneous hypersensitivity^3, 54, 58^. CGRP-induced sensitization of the trigeminal ganglia can propagate to third-order thalamic neurons and sensitization of these neurons can lead to extracephalic allodynia in chronic pain disorders^54^. There is an association between cutaneous allodynia and a higher headache frequency, suggesting its role as possible marker and risk factor for chronic migraine^59–61^. Additionally, the effective response of patients to migraine treatments, including triptans, appears to be reduced in patients with cutaneous allodynia^62, 63^. Upon administration of C21 in the hM4Di mice (inhibition of the bilateral PoT), we see a full rescue of CGRP-induced touch hypersensitivity, suggesting the PoT is a key sensory prosessing center for CGRP-induced migraine-like touch hypersensitivity in mice.

Photophobia is a common symptom of migraine that occurs when even a low level of ambient light is percieved as painful or uncomfortable and can exacerbate headache. The clinical manefestation of this symptom is widely believed to be a result of convergence of nociceptive signals from the SpV and signals from intrinsically photosensitive retinal ganglion cells in the PoT^35^. This is consistent with evidence suggesting that direct administration of CGRP into the PoT and optical stimulation of the PoT induces light aversion^37^. We found that inhibition of the PoT using C21-activation of hM4Di resulted in a partial rescue of the CGRP-induced light aversive phenotype in two strains of mice. The partial rescue of light aversion in both mouse strains suggest that other brain regions are involved in this phenotype alongside the PoT, for example the cerebellum^41, 47^. Our results emphasize the increasingly complex neurocircuitry of migraine given the distinct results for two sensory phenotypes that are surrogates for cutaneous allodynia and photophobia.

We repeated our experiments using C21 in a genetically diverse CD-1 strain of mice. Using multiple strains of mice ensures that the positive results we are observing are not restricted to a single mouse strain. The CD-1 results were consistent with the results from the C57BL/6J mice and help validate our conclusion that the PoT is necessary for touch hypersensitivity and partially necessary for light aversion in mice.

### Limitations

We used a total of four independent cohorts for this study. We were unable to conduct histology on one of the cohorts of C57BL/6J mice due to complications in the brain harvesting pipeline. While our mistargeted data suggest that hM4Di expression in the bilateral PoT, not the surrounding regions, is necessary for rescue of migraine-like behaviors, it is difficult to make strong conclusions from the underpowered small number of mistargeted mice. Similarly, it is difficult to determine whether unilateral expression of the DREADD in the PoT is sufficient to rescue the migraine-like phenotypes due to small sample size.

Further, the specific nuclei of the PoT that are responsible for the observed peripheral CGRP response remain to be determined. Additional studies with smaller injection volumes will be needed to specify nuclei within the PoT. While there are CGRP binding sites in the PoT ^37^, we think it is unlikely that peripheral CGRP is able to penetrate the blood brain barrier to act on the PoT directly^64^. The direct sites of CGRP action are still vigorously debated; more evidence is being reported in preclinical studies suggesting that both peripheral and central CGRP signaling underlie migraine phenotypes, with CGRP in the CNS likely contributing to the sensory abnormalities observed in migraine^11, 37, 41, 65, 66^. Our data suggest that the PoT could be a junction point for the actions of both central and peripheral CGRP actions. This supports the theory that peripheral and central neurocircuitry and signaling pathways are closely linked and both underlie migraine-like behaviors.

## Conclusion

In conclusion, this study indicates that the PoT in mice is necessary for migraine-like touch hypersensitivity, with partial necessity for light aversive behavior. These data highlight the PoT as a new central target for migraine therapeutics.

## Financial Support

This work was supported by the National Institutes of Health R01NS075599 (AFR) and VA merit award I01 RX003523 (LPS, AFR) and VA Center for the Prevention and Treatment of Visual Loss VA-ORD I50 RX003002 (LPS, AFR) from the United States (U.S.) Department of Veterans Affairs (Rehabilitation Research and Development Service).

## Abbreviations

CGRP: calcitonin gene-related peptide
CNS: central nervous system
ip: intraperitoneal
PoT: posterior thalamic nuclei
SpV: spinal trigeminal nucleus
DREADDs: Designer Receptors Exclusively Activated by Designer Drugs
hM4Di: modified form of the human M4 muscarinic receptor
C21: DREADD agonist compound 21
CNO: clozapine N-oxide
VA: Veterans Affairs
IACUC: Institutional Animal Care and Use Committee
ARRIVE: Animal Research: Reporting of *In Vivo* Experiments
AAV: adeno-associated virus
CaMKIIa: calmodulin kinase IIa
AP: anterior/posterior
ML: medial/lateral
DV: dorsal/ventral
PBS: phosphate buffered saline
TRITC: tetramethylrhodamine isothiocyanate
SEM: standard error of the mean
fMRI: functional magnetic resonance imaging
Tx: treatment

## Acknowledgements

pAAV-CaMKIIa-hM4D(Gi)-mCherry was a gift from Bryan Roth (Addgene plasmid # 50477; http://n2t.net/addgene:50477; RRID:Addgene_50477). Assay schematics were created in Biorender (A.M. Greenway (2025) https://BioRender.com/o66f767; https://BioRender.com/l61x847; https://BioRender.com/p53u934). Allen Mouse Brain Atlas reference images for histological confirmation of targeting were obtained from mouse.brain-map.org and atlas.brain-map.org.

## Notes

**<u>Conflict of Interest Statement:</u>** AFR serves as a consultant for Pfizer, Delphian Therapeutics, AbbVie, Lundbeck, and holds patents for uses of the CGRP monoclonal antibody, and the CGRP gene enhancer. LPS and AFR both hold patents for uses of the PACAP monoclonal antibody. The other authors declare no potential conflicts of interest with respect to the research, authorship, and/or publication of this article. The contents do not represent the views of the U.S. Department of Veterans Affairs or the United States Government.

### Competing Interest Statement

AFR serves as a consultant for Pfizer, Delphian Therapeutics, AbbVie, Lundbeck, and holds patents for uses of the CGRP monoclonal antibody, and the CGRP gene enhancer. LPS and AFR both hold patents for uses of the PACAP monoclonal antibody. The other authors declare no potential conflicts of interest with respect to the research, authorship, and/or publication of this article. The contents do not represent the views of the U.S. Department of Veterans Affairs or the United States Government.

