## Supplemental Figure 1 for "Inhibition of Posterior Thalamic Nuclei Attenuates CGRP-induced Migraine-like Behavior in Mice"

### SUPPLEMENTAL MATERIALS

**Supplemental Figure 1:** ***Histological mistargets for plantar von Frey assay using C21 in C57BL/6J and CD-1 mice suggest proper targeting is critical for DREADD effects.*** (**A** and **B**) mCherry control (A) and hM4Di experimental (B) C57BL/6J mice mistargets receiving 1X PBS or CGRP (0.1 mg/kg) treatment and C21 (1 mg/kg) + PBS or C21 + CGRP treatment. (**A**) CGRP and C21 + CGRP treatments do not induce a significant decrease in withdrawal threshold compared to PBS and C21 + PBS treatments, respectively, in the mCherry control mice. (**B**) CGRP successfully induces a significant decrease in withdrawal threshold compared to PBS treatment in hM4Di mice. C21 + CGRP administration in the hM4Di cohort does not alter withdrawal threshold compared to CGRP treatment. PBS and C21 + PBS treatments do not alter withdrawal threshold in neither mCherry nor hM4Di mice. (**C**) hM4Di experimental CD-1 mice mistargets receiving 1X PBS or CGRP (0.1 mg/kg) treatment and C21 (1 mg/kg) + PBS or C21 + CGRP treatment. CGRP fails to induce a significant decrease in withdrawal threshold in the hM4Di mice compared to PBS treatment, and this is unaffected by C21 administration. PBS and C21 + PBS treatments do not affect withdrawal threshold. Open and closed symbols denote males and females, respectively. Circle symbol denotes unilateral right PoT hit, triangle symbol denotes unilateral left PoT hit, and square symbol denotes complete miss of bilateral PoT. Data were transformed to enable statistical comparisons to be made. Mean +/- SEM 50% threshold shown. $*p \leq 0.05$, $**p \leq 0.01$. (**A** and **B**) Mistargeted mice from 2 independent cohorts of C57BL/6J mice. (**C**) Mistargeted mice from 1 independent cohort of CD-1 mice. For exact p-values and sample sizes, see Table 1.

**Supplemental Figure 1:**

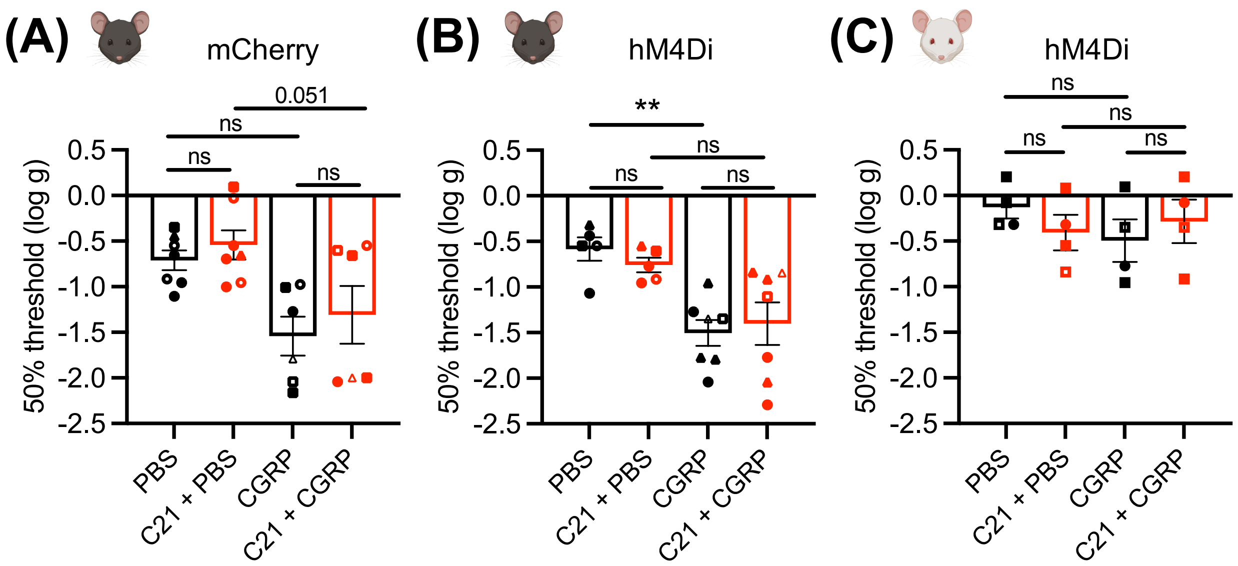
